# Discovery of anti-SARS-CoV-2 S2 protein antibody CV804 with broad-spectrum reactivity with various beta coronaviruses and analysis of its pharmacological properties in vitro and in vivo

**DOI:** 10.1101/2024.02.28.582480

**Authors:** Yoji Tsugawa, Kentaro Furukawa, Tomoko Ise, Masahiro Takayama, Takeshi Ota, Takayuki Kuroda, Shinya Shano, Takashi Hashimoto, Haruyo Konishi, Takeshi Ishihara, Masaaki Sato, Haruhiko Kamada, Keita Fukao, Takao Shishido, Tatsuya Takahashi, Satoshi Nagata

## Abstract

SARS-CoV-2 pandemic alerts us that spillovers of various animal coronaviruses to human in the future may bring us enormous damages. Thus, there is a significant need of antibody-based drugs to treat patients infected with previously unseen coronaviruses.CV804 against the S2 domain of the spike protein, which is less prone to mutations. CV804 shows not only broad cross-reactivities with representative 20 animal-origin coronaviruses but also with diseases-associated human beta coronaviruses including SARS-CoV, MERS-CoV, HCoV-OC43, HCoV-HKU1 and mutant strains of SARS-CoV-2. Other than that, the main characteristics of CV804 are that it has strong antibody-dependent cellular cytotoxicity (ADCC) activity to SARS-CoV2 spike protein-expressed cells in vitro and completely lacks virus-neutralization activity. Comprehensively in animal models, CV804 suppressed disease progression by SARS-CoV-2 infection. Structural studies using HDX-MS and point mutations of recombinant spike proteins revealed that CV804 binds to a unique epitope within the highly conserved S2 domain of the spike proteins of various coronaviruses. Based on the overall data, we suggest that the non-neutralizing CV804 antibody recognizes the conformational structure of the spike protein expressed on the surface of the infected cells and weakens the viral virulence by supporting host immune cells’ attack through ADCC activity in vivo. CV804 epitope identified in this study is not only useful for the design of pan-corona antibody therapeutics but also to design next-generation coronavirus vaccines and antiviral drugs.

## Introduction

The COVID-19 pandemic caused by SARS-CoV-2 beta coronavirus discovered in 2019 has had a severe impact on the global economy and health^1^. Besides COVID-19, various corona viruses caused various infectious diseases likely by animal-to-human spillover infection and pose a significant threat to public health. The coronaviridae is classified into four genera based on antigenicity and genetic criteria: alpha, beta, gamma, and delta^2^. Common human coronaviruses, such as HCoV-OC43, HCoV-HKU1 (beta coronaviruses), and HCoV-229E and HCoV-NL63 (alpha coronaviruses), are known to circulate annually and cause mild to moderate upper respiratory tract diseases^3,4^. In addition, the severe acute respiratory syndrome coronavirus (SARS-CoV) that occurred from 2002 to 2003 and the Middle East respiratory syndrome coronavirus (MERS-CoV) that occurred in 2012 are representative examples of highly pathogenic coronaviruses that caused pandemics with high mortality rates^5^. These historical data strongly suggest that zoonotic spillover of corona viruses will likely occur in the future^6^. Therefore, to prepare possible next pandemic, making the development of broad-spectrum antiviral drugs and vaccines against coronaviruses is a pressing need for proactive measures against coronavirus.

Anti-virus spike antibodies show promise as effective drugs capable of promptly addressing new coronavirus-derived infections in case of future pandemics; however, the high specificities prevent their common use to treat different coronaviruses and easily allow virus-mutation to escape from the antibodies^7^. So far, virus neutralization, preventing virus entry into the host cells, is the most widely pursued mechanism for antibody drugs. The target portion of the most of the neutralizing antibodies (nAbs) that have been reported against SARS-CoV-2, SARS-CoV-1, and MERS-CoV is the receptor-binding domain (RBD) in the S1 domain, which inhibit the virus from binding to target cells. However, obtaining anti-RBD antibodies with broad reactivity against diverse viruses and overcoming the escape mutations pose difficulties ^8–12^. On the other hand, The S2 domain shows a higher degree of conservation across various coronaviruses than the S1 domain. Although the expected wide cross-reactivity is attractive for developing antibody drugs to the S2 domain for many corona viruses, neutralization limits the targetable antibody epitope regions in few regions of the S2 domain. Indeed, existing antibodies targeting S2, such as B6 and S2P6, binds to the restricted membrane proximal regions and work by preventing the refolding of the S2 subunit and inhibiting membrane fusion, that is essential event for the virus to introduce the viral genome into the host cell cytoplasm to replicate itself^13–16^. Antibodies to another S2 epitope such as 76E1 have also been reported to inhibit S2’ cleavage and membrane fusion, thereby demonstrating neutralizing activity^17^.

Antibodies targeting a less variable region outside the RBD, specifically the S2 domain in the virus spike, have not undergone extensive validation as pharmaceuticals due to their limited in vitro infection-blocking activity (virus neutralization). Nonetheless, even antibodies lacking neutralizing activity that were induced in infected patients may support the host immune mechanisms, such as antibody-dependent cellular cytotoxicity (ADCC), contributing to favorable patient outcomes and preventing infection from related viruses^18,19^.

Therefore, we expanded and explored targetable regions for therapeutic antibodies in less mutable S2 domain in this study. We hypothesized that the intrinsic ADCC function of antibody is significant to exhibit the therapeutical capability even without virus-neutralization.

We successfully identified a cross-reactive antibody called CV804 against representative strains of beta coronaviruses. This antibody, CV804, inhibited the exacerbation of disease caused by viral infection through effector activity since it does not possess neutralizing activity. Structural and functional analysis revealed that CV804 targets a unique epitope within the highly conserved S2 domain of the spike protein. Identifying highly conserved epitopes holds promise for the design of broad-spectrum coronavirus vaccines and antiviral agents against current and future emerging SARS-CoV-2 mutants and other coronavirus genera.

## Materials and Methods

### Cells and viruses

VeroE6/TMPRSS2 cells from the National Institutes of Biomedical Innovation (Tokyo, Japan) were used to evaluate the antiviral activity against SARS-CoV-2. Cells were maintained in Dulbecco’s modified Eagle’s medium (Thermo Fisher Scientific) supplemented with 10% heat-inactivated fetal bovine serum (FBS) at 37 °C with 5% CO2. ExpiCHO cells (Thermo Fisher Scientific) were maintained in ExpiCHO expression medium (Thermo Fisher Scientific) at 37 °C under 8% CO2. Expi293F cells (Thermo Fisher Scientific) were maintained in Expi293 expression medium (Thermo Fisher Scientific) at 37 °C under 8% CO2.

SARS-CoV-2 clinical isolates were obtained from the National Institute of Infectious Diseases (NIID; Tokyo, Japan): hCoV-19/Japan/TY/WK-521/2020 (Pango Lineage: A), hCoV-19/Japan/QK002/2020 (B.1.1.7), hCoV-19/Japan/QHN001/2020 (B.1.1.7), hCoV-19/Japan/QHN002/2020 (B.1.1.7), hCoV-19/Japan/TY7-501/2021 (P.1), hCoV-19/Japan/TY8-612/2021 (B.1.351), hCoV-19/Japan/TY11-927/2021 (AY.122), hCoV-19/Japan/TY33-456/2021 (C.37), hCoV-19/ Japan/TY28-444/2021 (P.3), hCoV-19/ Japan/TY38-873/2021 (BA.1.18), hCoV-19/ Japan/TY38-871/2021 (BA.1.1), hCoV-19/ Japan/TY40-385/2022 (BA.2), hCoV-19/ Japan/TY41-721/2022 (BA.2.12.1), hCoV-19/ Japan/TY41-716/2022 (BA.2.75), hCoV-19/ Japan/TY41-703/2022 (BA.4.1), hCoV-19/ Japan/TY41-763/2022 (BA.4.6), hCoV-19/Japan/TY41-702/2022 (BE.1), hCoV-19/ Japan/TY41-704/2022 (BA.5.2.1), hCoV-19/ Japan/TY41-796/2022 (BQ.1.1), hCoV-19/ Japan/TY41-795/2022 (XBB.1), hCoV-19/ Japan/TY41-686/2022 (XE), hCoV-19/ Japan/TY41-820/2022 (BF.7), hCoV-19/Japan/TY41-828/2022 (BF.7.4.1), hCoV-19/ Japan/23-018/2022 (XBB.1.5), and hCoV-19/Japan/TY7-503/2011 (P.1). SARS-CoV-2 strains were propagated in VeroE6/TMPRSS2 cells, and infectious titers were determined using the standard tissue culture infectious dose (TCID)50 method in VeroE6/TMPRSS2 cells.

### Animal experiments and approvals

Animal studies were conducted under applicable laws and guidelines. The protocols received approval from the Shionogi Pharmaceutical Research Centre Institute Director (Shionogi & Co., Ltd., Toyonaka, Japan) based on the report of the Institutional Animal Care and Use Committee. (Assurance number: S21043D). All animals were kept in specific pathogen-free conditions, under regulated temperature and a 12-hour light and dark cycle. They had access to water and standard laboratory chow ad libitum, and the humidity was maintained at 30-70%. Virus inoculations were performed under anesthesia, with measures taken to minimize animal suffering. The in vivo studies were not conducted in a blinded manner, and animals were randomly assigned to infection groups.

Sample size calculations were not performed for each study; instead, sample sizes were determined based on previous in vivo virus challenge experiments. Twelve-week-old female BALB/c mice (n = 5) were inoculated via the intranasal route with 1.0×10^5^ TCID_50_/mouse (in 50 μl) of SARS-CoV-2 hCoV-19/Japan/TY7-501/2021 (gamma strain). Immediately after infection, the mice were intravenously administered 40 mg/kg of CV804, CV804 (LALA), REGN10987 or isotype control mAb (C1.18.4). As a reference, PBS-inoculated uninfected control mice were also prepared. Body weight and survival were monitored once daily until day 6 post-virus infection. At 6 days post-infection, the mice were euthanized. If the mice lost more than 20% of their body weight compared to their initial body weight according to humane endpoints, they were immediately euthanized and regarded to be dead in the analysis for survival time. The numbers of mice that survived, euthanized according to humane endpoints, or died before reaching the humane endpoints are summarized in S1 Table.

### SARS-CoV-2 spike IgG ELISA

In brief, the SARS-CoV-2 spike IgG ELISA is an indirect ELISA that is based on antibody/antigen interactions. Mouse antibodies were added to a solid-phase plate coated with anti-mouse IgG antibodies and incubated overnight at 4°C. After three washes with ELISA wash buffer, S2 protein (Acro BIOSYSTEMS) and trimeric S2 protein, along with S protein, were added and incubated for 1 hour at room temperature. Following three washes with ELISA wash buffer, the plate was incubated with anti-histidine tag antibody for 1 hour at room temperature. After another three washes with ELISA wash buffer, alkaline phosphatase-labeled anti-human IgG antibody was added and incubated for 1 hour at room temperature. Following four washes with ELISA wash buffer, color development was achieved by adding P-nitrophenyl phosphate solution, and absorbance was measured at 405 nm.

### Establishment of the in-cell SARS-CoV-2 ELISA

To detect SARS-CoV-2 infection, cells were distributed at a concentration of 3x10^5^ cells/ml. The culture medium was replaced, and a concentrated virus solution was introduced at 100 μL/well. The cells were then incubated at 37°C for 1 hour. The virus-infected medium was aspirated and substituted with a mixture of 2xMEM/8% FBS medium and 2% MC in a 1:1 ratio. The medium was subsequently exchanged with 1xMEM/4% FBS/1%MC at 200 μL/well. The cells were incubated at 37°C for 16-17 hours. The medium was gently removed, leaving approximately 30 μL in each well. A single wash with PBS was performed, ensuring complete removal of the PBS. A 4% paraformaldehyde solution was added at 300 μL/well and allowed to incubate at room temperature for 30 minutes. The plate was then wiped with ethanol, retaining about 30 μL, before removal of the 4% paraformaldehyde. The cells were washed twice with PBS. A single wash with wash buffer was performed, leaving behind approximately 50 μL before removal. A 2x Permeability buffer was introduced at 50 μL/well and left at room temperature for 10 minutes. Subsequently, the cells were washed three times with wash buffer, retaining 50 μL after each wash. A 2% BSA solution was added at 50 μL/well and allowed to incubate at room temperature for 30 minutes. The blocking solution was completely aspirated, and the primary antibody was added at 50 μL/well. The cells were incubated at room temperature for 2 hours. Secondary antibody: The cells were washed four times with 300 μL of wash buffer. The wash buffer was completely removed, and 50 μL of secondary antibody, diluted 2000-fold in 0.1% BSA solution, was added to each well. The cells were incubated at room temperature for 1 hour. The cells were washed four times with 300 μL of wash buffer. After three washes, staining was visualized by addition of TrueBlue detection reagent. Infected foci were then enumerated using Immunospot (Cellular Technology Limited).

### Flow cytometry

The antibody-reacted cells were analyzed by flow cytometry (FCM). Plasmids expressing spike proteins from various coronaviruses, including SARS-CoV-2, its strains (alpha, delta, lambda, theta, mu, omicron BA1, BA2, BA5, BQ1.1, XBB, XE), SARS-CoV-1, MERS, Human coronaviruses 229E, HKU1, NL63, OC43, Bat-CoV, Bat-HpCoV, Bat-HKU9, Bat-HKU4, Bovine-L9, MHV, PCoV GX-P1E, Bat-HKU3, BtRs-CoV, SADS-CoV, ChRCoV HKU24, Ro-BatCoV, LongquanAa mCoV, Eidolon BatCoV, Chaerephon BatCoV, RodentCoV, BtVs-BetaCoV, Rhinolophus BatCoV HKU, NL63-related BatCoV, WigeonCoV HKU20, BulbulCoV HKU11-934, PorcineCoV HKU15, MuniaCoV HKU13-3514, Beluga whaleCoV SW1 were constructed and transfected into HEK293 cells. The transfected cell populations were separately allowed to react with the obtained antibodies at 4°C for 30 minutes. EC_50_ values were determined by plotting the compound concentration versus inhibition and fitting data with a four-parameter logistical fit (Model 205, XLfit).

### Neutralizing antiviral activity

Antiviral activities against SARS-CoV-2 were evaluated using VeroE6/TMPRSS2 cells. The viral strains hCoV-19/Japan/TY/WK-521/2020 and SARS-CoV-2/Japan/WK-521/2020 were employed, utilizing VeroE6/TMPRSS2 cells as the infected cells. The culture medium consisted of MEM supplemented with 2% FBS. For antibody dilution, CV804, REGN10987, Ly-CoV1404 and isotype control antibody were adjusted to a maximum concentration of 3.3 μM, each antibody with a 3-fold serial dilution of 10 points. The virus was adjusted to a concentration of 3000 TCID_50_/well. The adjusted antibody dilutions and virus solution were mixed in equal proportions and allowed to react at room temperature for 1 hour. The virus-antibody mixture was then added to VeroE6/TMPRSS2 cells at a density of 15,000 cells/well and cultured at 37℃, 5% CO2 conditions. After 72 hours, CellTiter-Glo 2.0 (Promega) was added to measure the fluorescence intensity. Antiviral activity against SARS-CoV-2 was assessed by monitoring the cell viability. EC_50_ values were determined by plotting the compound concentration versus inhibition and fitting data with a four-parameter logistical fit (Model 205, XLfit).

### Measurement of ADCC activity

HEK293 cells expressing spike protein on their cell membrane were cultured overnight. Mouse antibodies (CV801, CV804, CV820, CV925, CV1117) and effector cells (Promega) expressing mouse FcγRIV and NFAT luciferase reporter were added to the cells, and the reaction was conducted at 37°C in a 5% CO2 incubator for 6 hours. Next, a colorimetric reagent was added and allowed to react for 5 minutes, followed by detection of the fluorescent signal. Four out of the five antibodies, except for CV925, were found to have ADCC activity. The EC_50_ values for each antibody indicate that the ADCC activity was below the detectable limit and therefore the value could not be determined.

### HDX-MS analysis

The plasmids for the recombinant SARS-CoV-2 variants were transiently transfected into Expi293F cells (Thermo Fisher Scientific) with the ExpiFectamine 293 Transfection Kit (Thermo Fisher Scientific) according to the manufacturer’s protocol. After 4–5 days of culture in the Expi293 Expression Medium (Thermo Fisher Scientific), supernatants were collected and passed through a 0.22-µm filter. The recombinant proteins were purified from supernatants by NGL COVID-19 Spike Protein Affinity Resin (Repligen) according to the manufacturer’s protocol. The elute with 100 mM glycine-HCl (pH 2.7) from the resin was immediately neutralized with 1/10 faction volume of 1 M Tris-HCl pH 8.5 and concentrated using an Amicon Ultra-2 Centrifugal Filter Unit 100kDa NWMCO (Merck Millipore), in which the elution buffer was exchanged with PBS (GIBCO).

The hydrogen/deuterium exchange method has been previously described^29,30^. The system for HDX-MS experiments used the HDX-PAL system for deuterium labeling (Leap Technologies Inc.), enzyme pepsin column for sample digestion (Waters), HyperSil Gold column as a trap column (Thermo Fisher Scientific), Accelaim PepMap300 C18 column as an analytical column (Thermo Fisher Scientific), and Orbitrap Eclipse for mass measurement of digested peptides (Thermo Fisher Scientific). The target protein for hydrogen/deuterium exchange epitope mapping was a chimeric protein of the spike protein of SARS-CoV-2 expressed in HEK293 cells. The chimeric protein used included the ectodomain of the spike protein of SARS-CoV-2 (amino acids 1-1213 of the S protein described in Genbank ACC No. QHD43416.1), with amino acids 986 (K) and 987 (V) replaced with proline (P). The deuterated buffer was prepared at pH 7.4 using 10 mM PBS buffer adjusted with D2O. The above target protein was allowed to react at 37°C for 30 minutes in the presence or absence antibody. The initiation of deuterium labeling, control of reaction time, quenching reaction, injection into the UPLC system, and control of digestion time were all carried out automatically by the HDX-PAL system. First, the target protein/antibody complex or target protein was diluted 10-fold with deuterated buffer to initiate deuterium labeling. Deuterium labeling was carried out under conditions of 10°C for 30, 60, 120, and 240 seconds of labeling time. The deuterium-labeled sample (27 μL) was mixed with an equal volume of quenching buffer (4 mol/L guanidine hydrochloride, 0.2 mol/L glycine hydrochloride, 0.5 mol/L TCEP) and quenched at 0°C for 3 minutes. Subsequently, the quenched sample (50 μL) was injected into an Enzyme Pepsin Column for online pepsin digestion. The digested peptides were trapped on a trap column (Hypersil Gold column) at 1°C and eluted onto an analytical column (Acclaim PepMap300 C18) using a 7-minute gradient separation of 10%-35% B (mobile phase A: 0.1% formic acid in water, mobile phase B: 0.1% formic acid/acetonitrile). Mass spectrometry (Orbitrap Eclips) was set with an electrospray voltage of 4000 V, a scan time of 1 second, and a mass/charge range of 260-2000 in positive ion mode to acquire both full-scan and MS/MS spectra. To identify the peptides from the acquired mass spectra, a search was conducted against a database containing the amino acid sequences of the spike protein of SARS-CoV-2 using Byos software (Protein Metrics). The retention time and mass information of the identified peptides were then imported into HDExaminer software (Sierra Analytics) to automatically calculate the deuterium exchange rates of each peptide in the hydrogen/deuterium exchange experiment. Peptides with calculated deuterium exchange rates were filtered according to the criterion that the difference in deuterium exchange rates (ΔD%) between the antibody present and absent samples of two or more adjacent peptides was 10% or more. The amino acid residues of the spike protein of SARS-CoV-2 corresponding to the region containing the peptides meeting the criterion were identified as the epitope of the antibody.

## Results

### Antibody isolation and binding activity to S protein

To obtain antibodies that effectively bind to a wide range of coronaviruses, we immunized mice with various immunization methods and their combinations using plasmid DNA encoding the S2 subunit of the spike protein of SARS-CoV2, proteins, spike-expressing cells, peptides. Using the antiserum, we confirmed their binding to the S2 region and coronaviruses, including MERS and SARS. Hybridomas were generated using the PEG and electrofusion methods, and specific clones with binding activity to HEK293 cells expressing the spike protein of SARS-CoV-2 were selected. We finally obtained five antibodies (CV801, CV804, CV820, CV925, CV1117). Their binding activities against different high-ordered structures of the spike protein were examined in ELISA using S2 protein monomers, trimers, and full-length S protein trimers (Fig 1).

**Fig 1.**
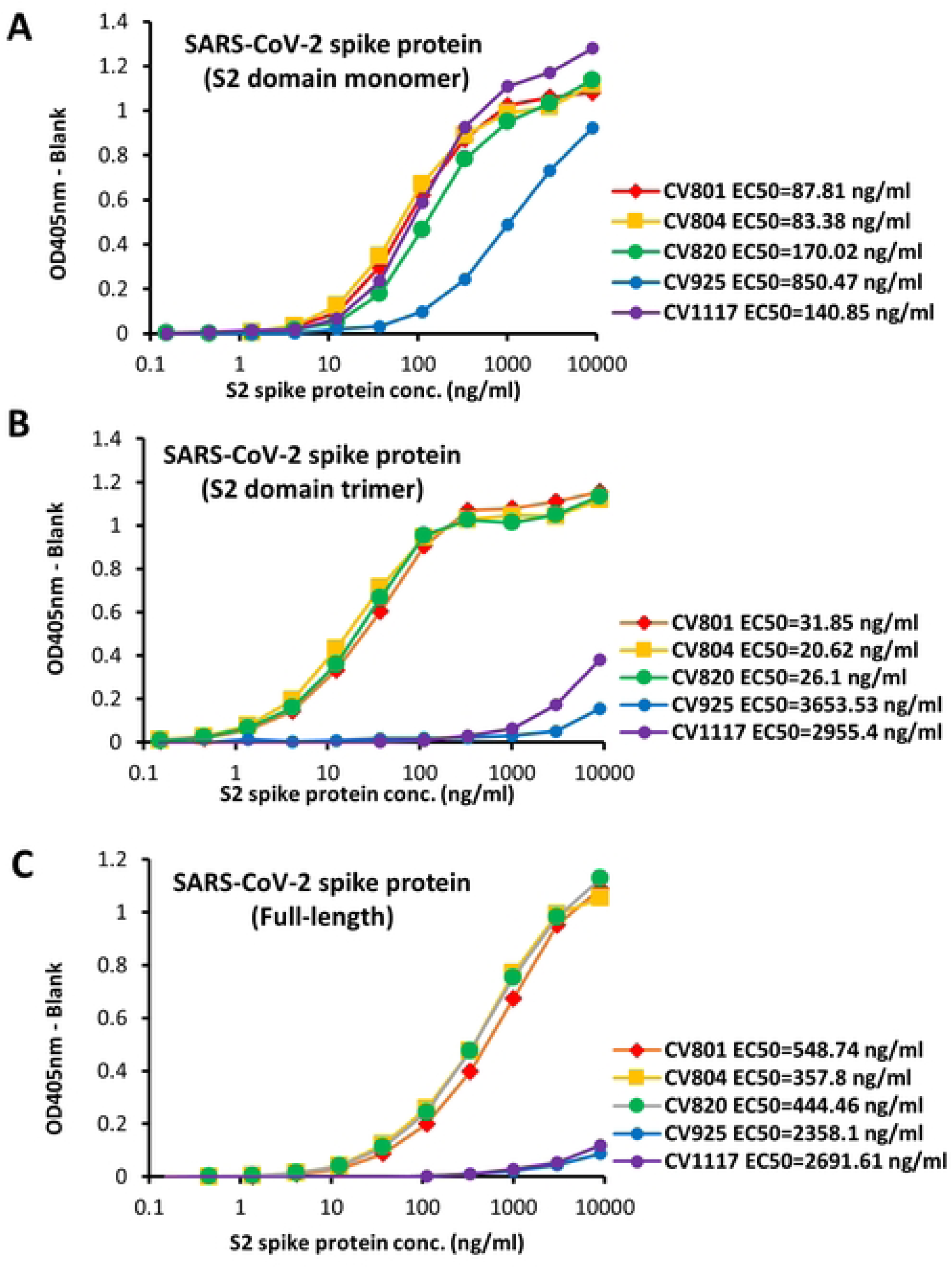
Antibody isolation and binding activity to S protein. 5 antibodies (CV801, CV804, CV820, CV925, CV1117) binding to the soluble S2 domain monomer (A), trimer (B), and full-length (C) of SARS-CoV-2 spike protein was assessed using ELISA assay. Representative data of two independent experiments are shown (A-C).

Although remarkably, all proteins demonstrated binding activity (Fig 1A). Interestingly, three antibodies (CV801, CV804, and CV820) displayed stronger affinity for the trimer compared to the monomer (Figs 1A-1C). To explore the binding activity of each antibody in greater detail, we examined their efficacy against single mutations and mutant strains of the S2 protein, which have been identified globally. Plasmids expressing various single mutants and mutant strains were created, transiently expressed in cells, and subjected to flow cytometry analysis. Interestingly, CV804 demonstrated exceptional effectiveness against confirmed single mutants from around the world, displaying robust binding activity (Table 1) (S1A Fig). Furthermore, to assess the cross-reactivity of this antibody, flow cytometry was performed on cells expressing S proteins from different types of coronaviruses. The results revealed that CV804 exhibited binding activity against a broad spectrum of coronaviruses, including SARS-COV2, SARS-COV1, MERS, Human-Coronavirus HKU1, NL63, OC43, Bat-COV RaTG13, Bat-HpCOV, Bat-HKU4, PCOV GX-P1E, Bat-HKU3-3, BtRs-COV, and SARS-COV (Table 2) as well as ChRCoV HKU24, Ro-BatCoV, LongquanAa mCoV, Eidolon BatCoV, Chaerephon BatCoV, RodentCoV, BtVs-BetaCoV, Rhinolophus BatCoV HKU, NL63-related BatCoV, WigeonCoV HKU20, BulbulCoV HKU11-934, PorcineCoV HKU15, MuniaCoV HKU13-3514, Beluga whaleCoV SW1 (S1B Fig).

**Table 1.**
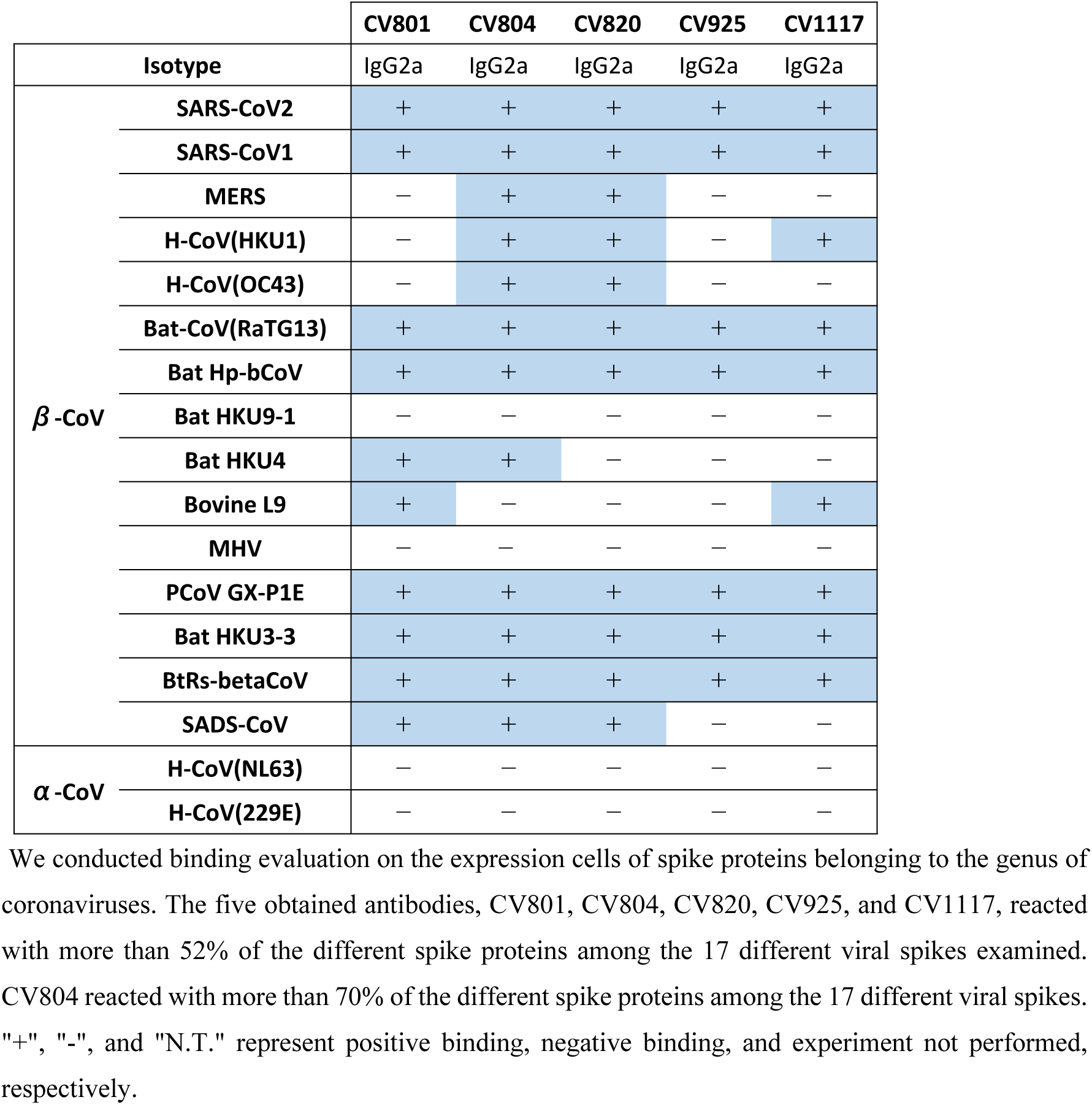
The presence or absence of binding to each viral strain is indicated.

|  |  | CV801 | CV804 | CV820 | CV925 | CV1117 |
| --- | --- | --- | --- | --- | --- | --- |
| Isotype |  | IgG2a | IgG2a | IgG2a | IgG2a | IgG2a |
| **β-CoV** | **SARS-CoV2** | + | + | + | + | + |
|  | **SARS-CoV1** | + | + | + | + | + |
|  | **MERS** | — | + | + | — | — |
|  | **H-CoV(HKU1)** | — | + | + | — | + |
|  | **H-CoV(OC43)** | — | + | + | — | — |
|  | **Bat-CoV(RaTG13)** | + | + | + | + | + |
|  | **Bat Hp-bCoV** | + | + | + | + | + |
|  | **Bat HKU9-1** | — | — | — | — | — |
|  | **Bat HKU4** | + | + | — | — | — |
|  | **Bovine L9** | + | — | — | — | + |
|  | **MHV** | — | — | — | — | — |
|  | **PCoV GX-P1E** | + | + | + | + | + |
|  | **Bat HKU3-3** | + | + | + | + | + |
|  | **BtRs-betaCoV** | + | + | + | + | + |
|  | **SADS-CoV** | + | + | + | — | — |
| **α-CoV** | **H-CoV(NL63)** | — | — | — | — | — |
|  | **H-CoV(229E)** | — | — | — | — | — |

**Table 2.**
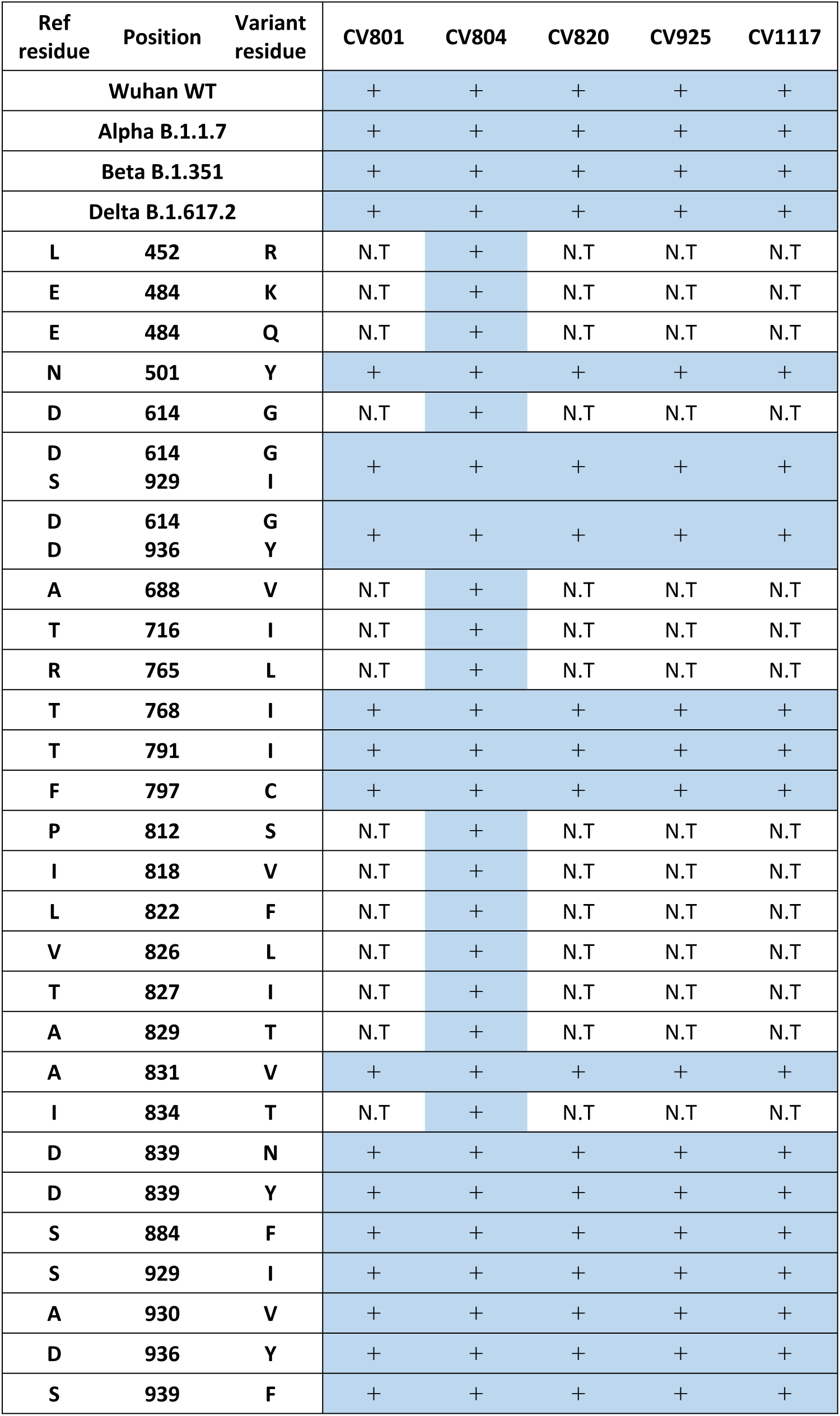
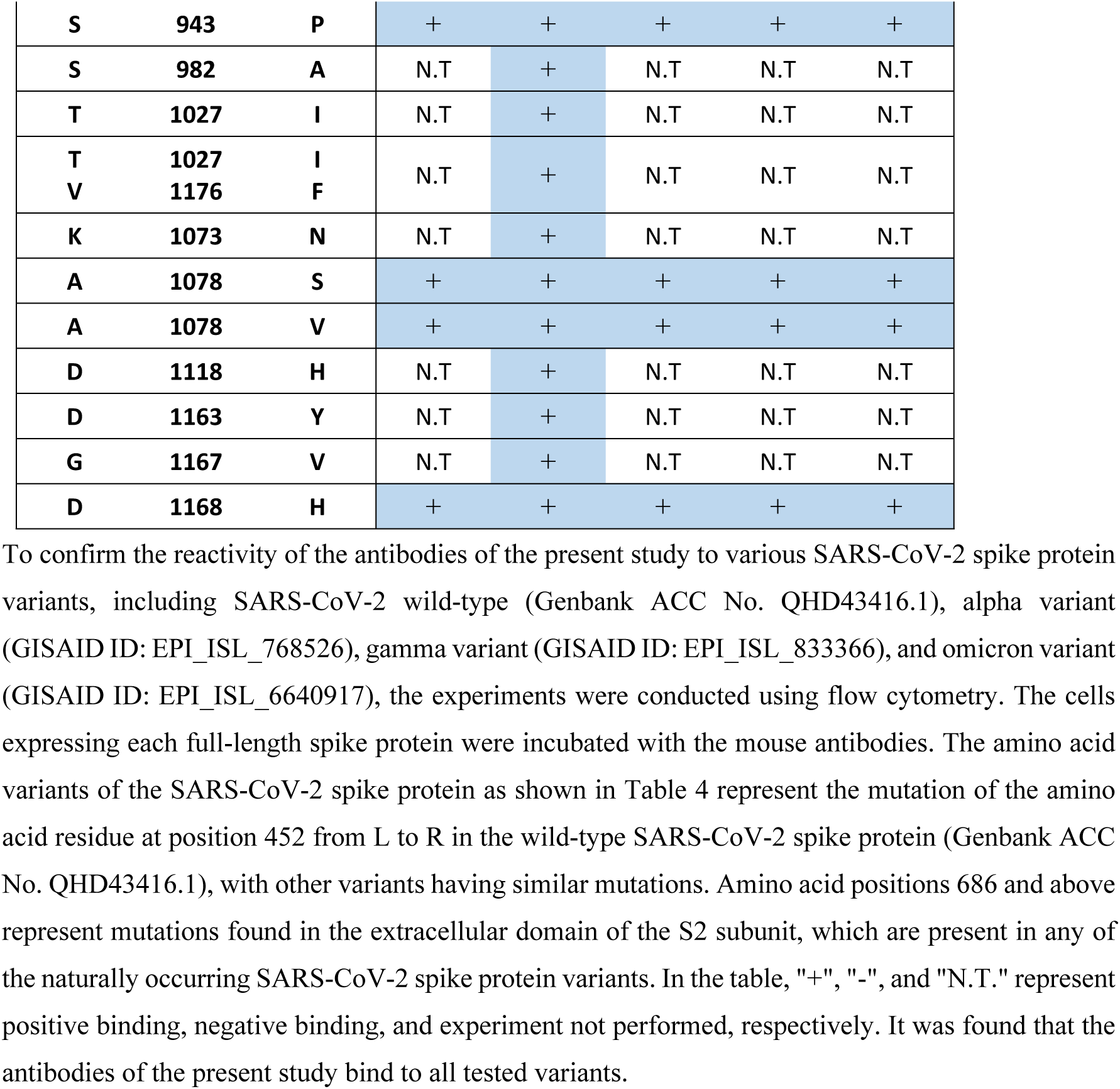
Reactivity to SARS-CoV-2 spike protein variants expressed in cells.

**Table 3.**
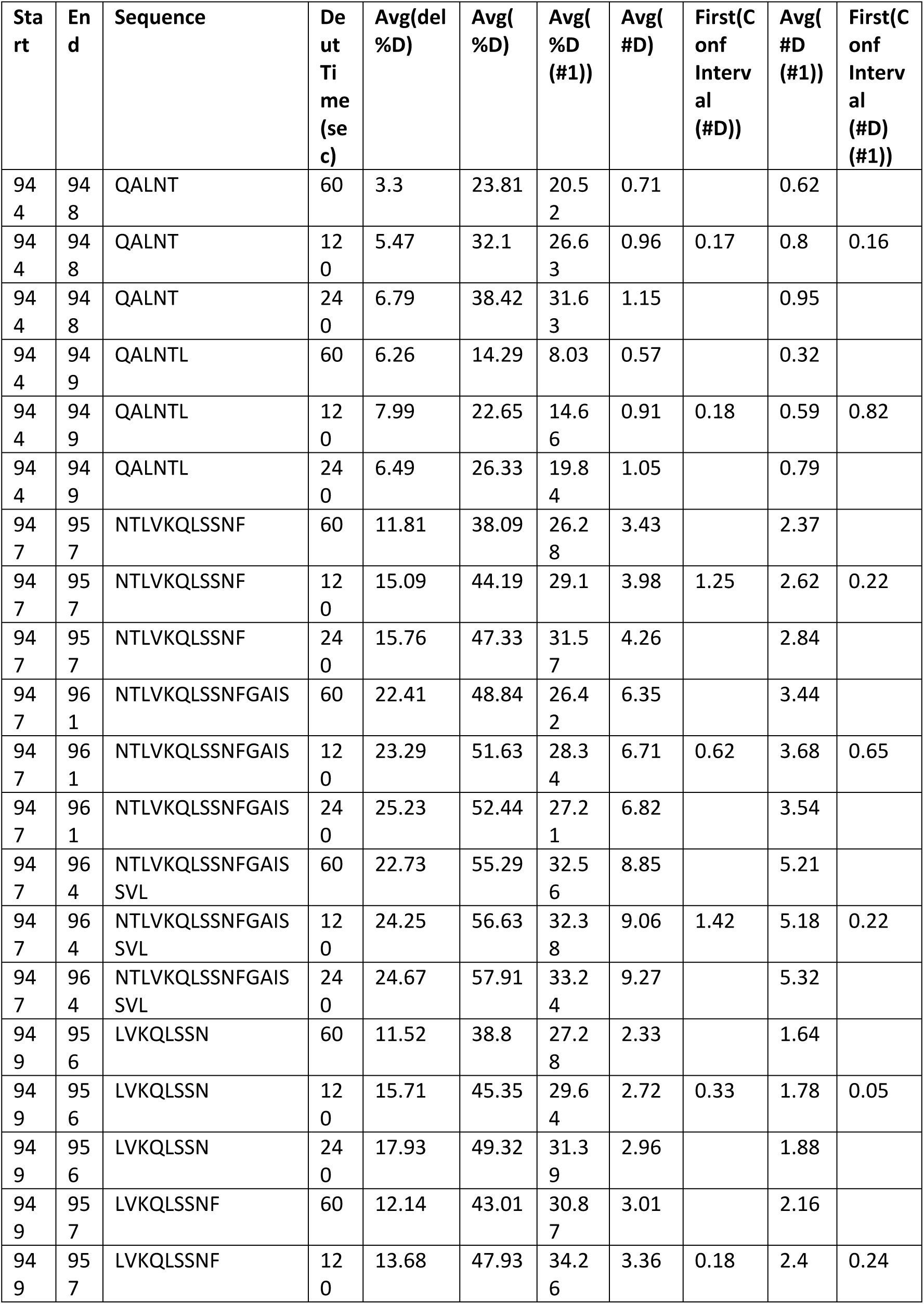
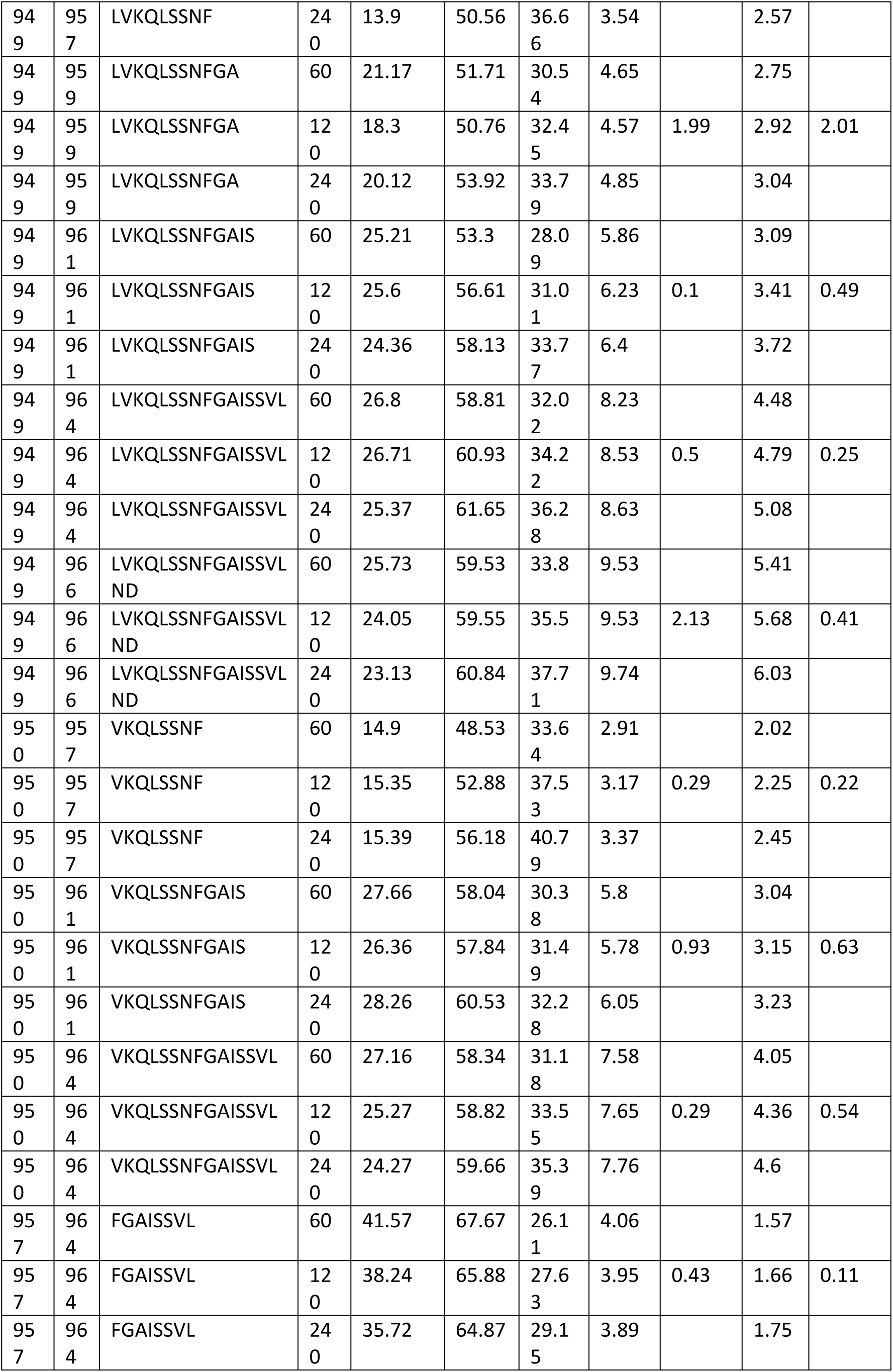
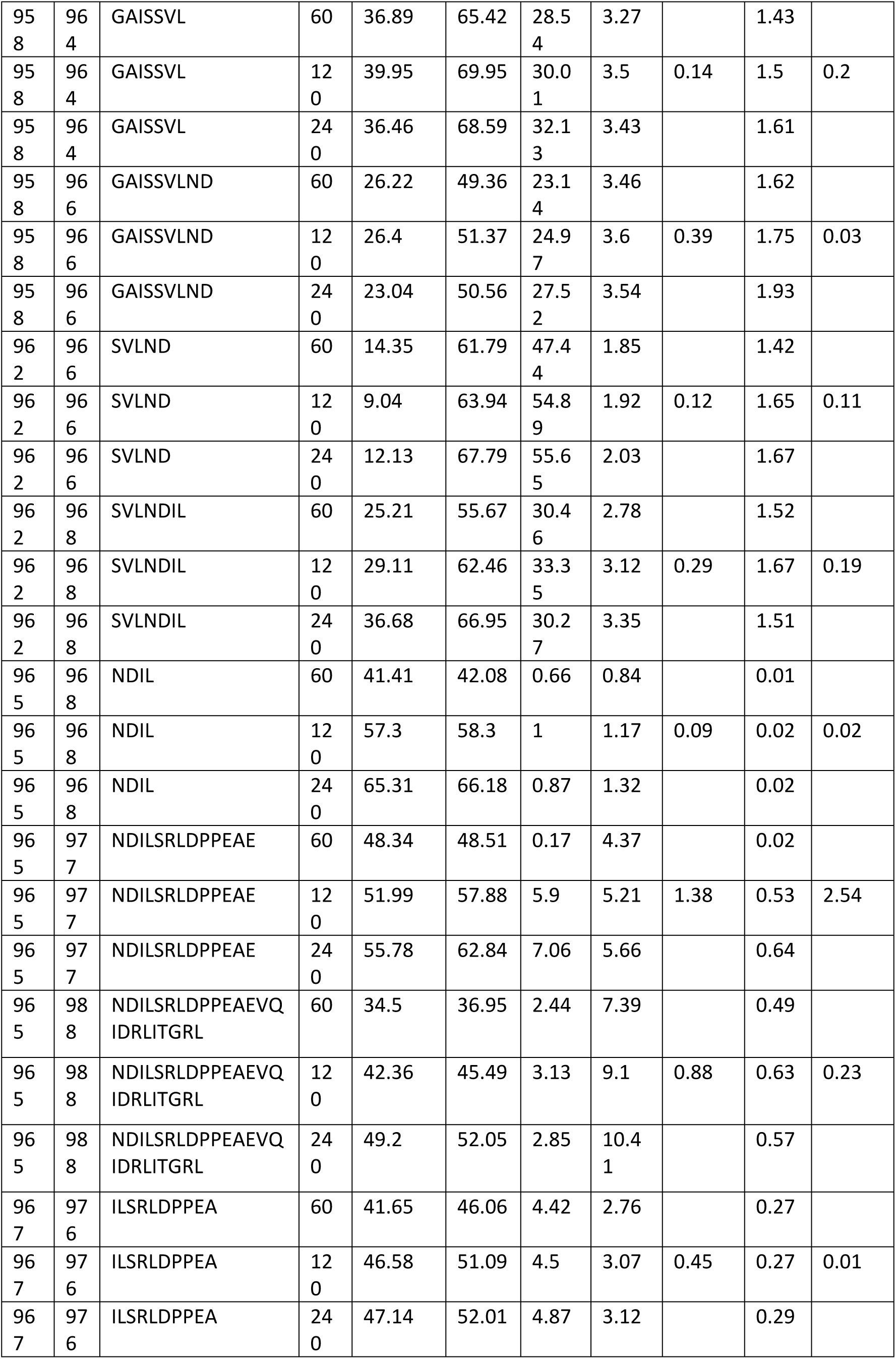
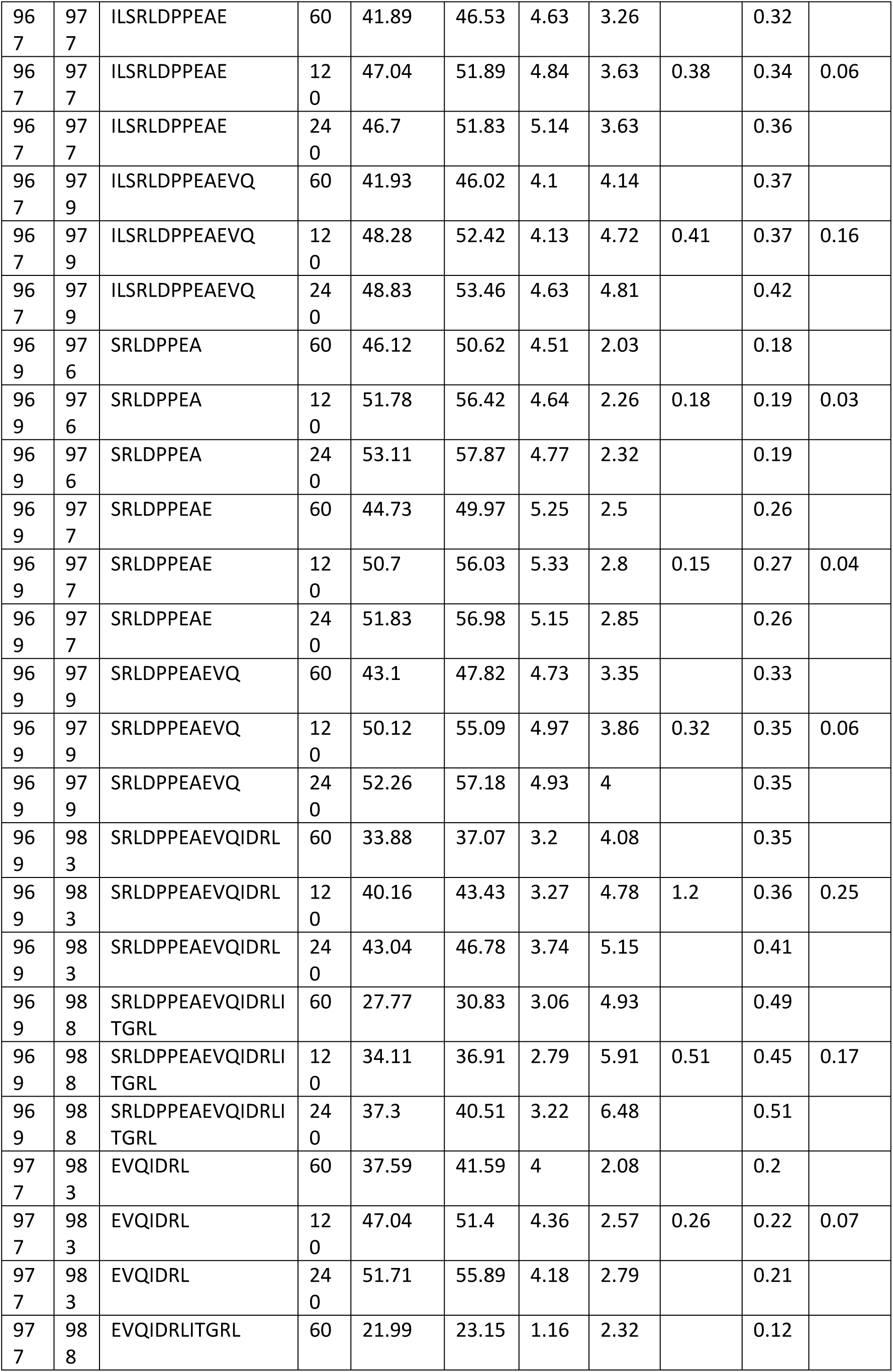
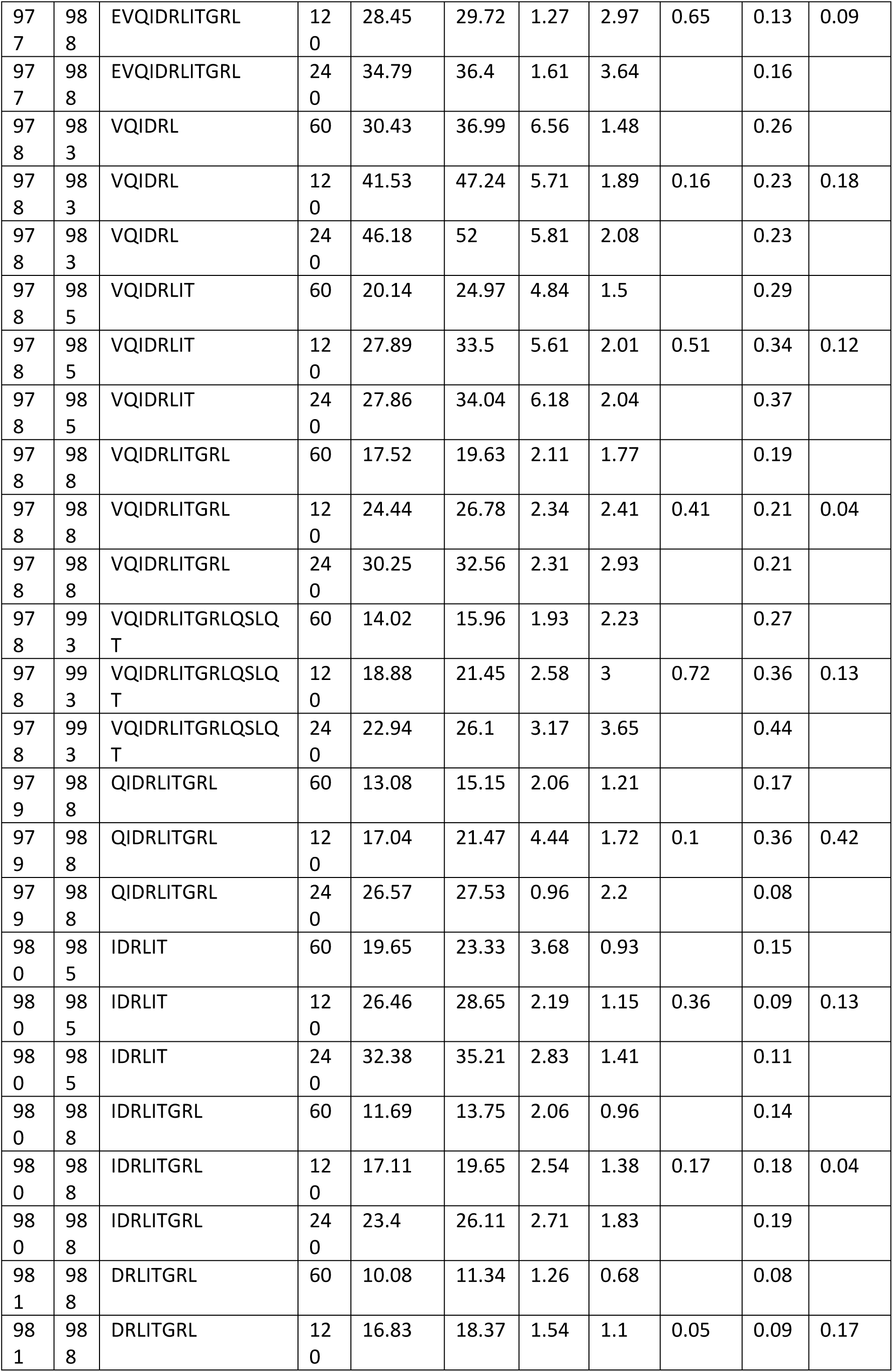
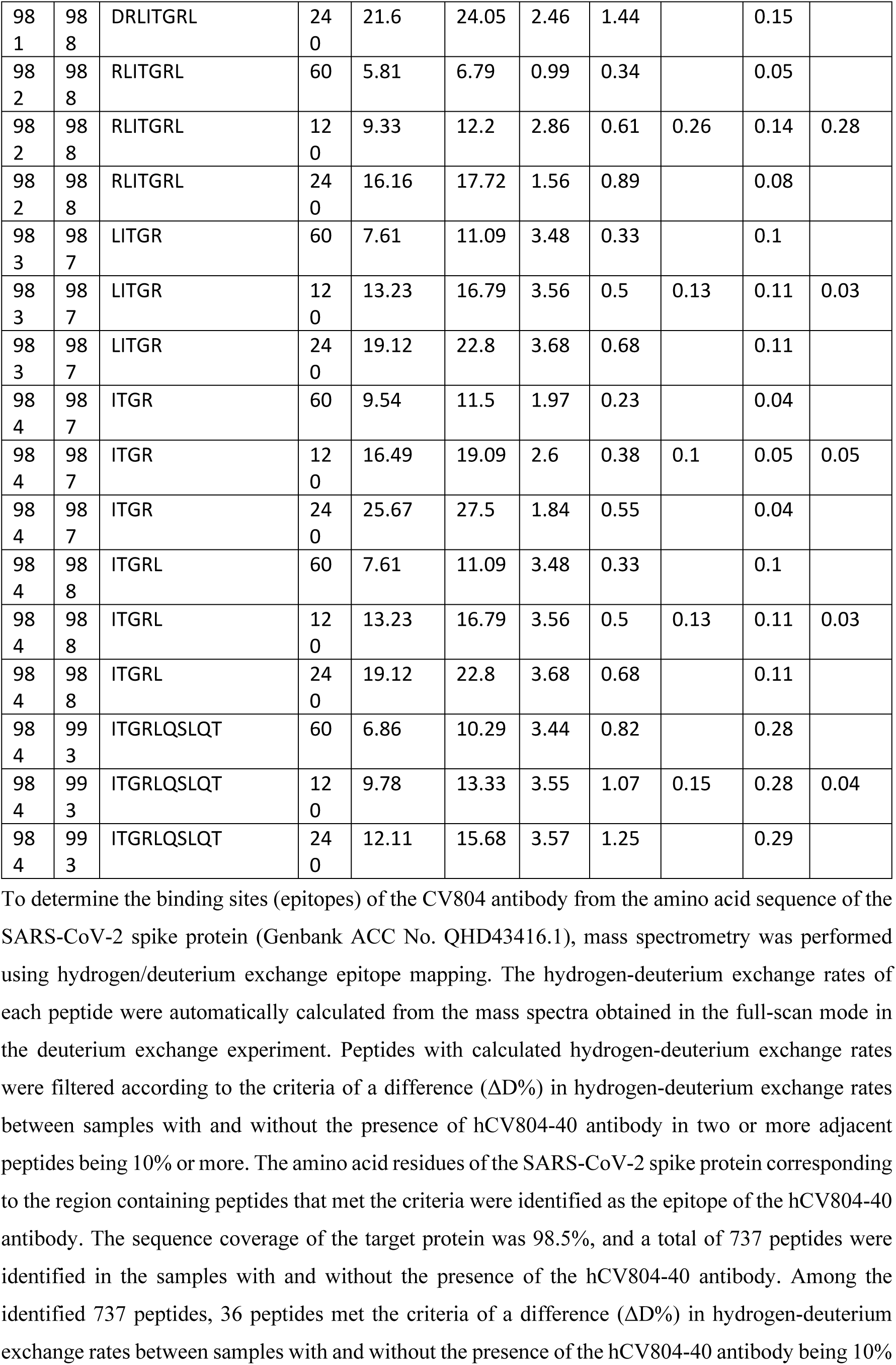
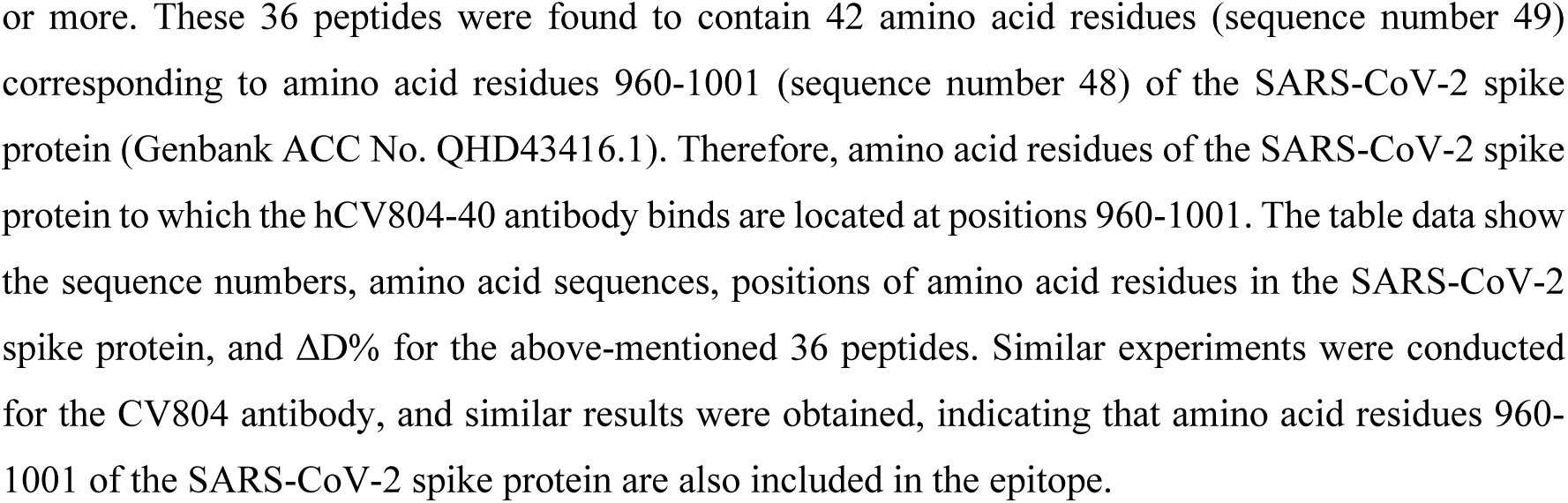
Epitope mapping of CV804 antibody binding to the SARS-CoV-2 spike protein using the HDX method.

### CV804 binds to SARS-CoV-2 and its variants but does not neutralize them

CV804, located within the S2 domain of the viral spike protein, exhibits a structure that is resistant to mutations, allowing it to effectively react with various mutant strains, including the Omicron strain, as well as numerous related coronaviruses^20^. However, there are still questions regarding the binding mechanism and functional characteristics of CV804.

For therapeutic purposes in humans, the mouse antibody CV804 (IgG-κ) was humanized using our proprietary algorithm (Shionogi Co. Ltd., Osaka, Japan). The variable regions were humanized based on the Kabat antibody numbering scheme, with substitutions of amino acids guided by physical property predictions from the closest germline and known human antibody sequences and structures. Consequently, the humanized antibody named hCV804 was generated. This antibody was produced and assessed for binding to the S protein. Remarkably, the humanized antibody maintained binding activity to the S protein, with a similar Kd value to that of the mouse-derived CV804 (Fig 2A).

**Fig 2.**
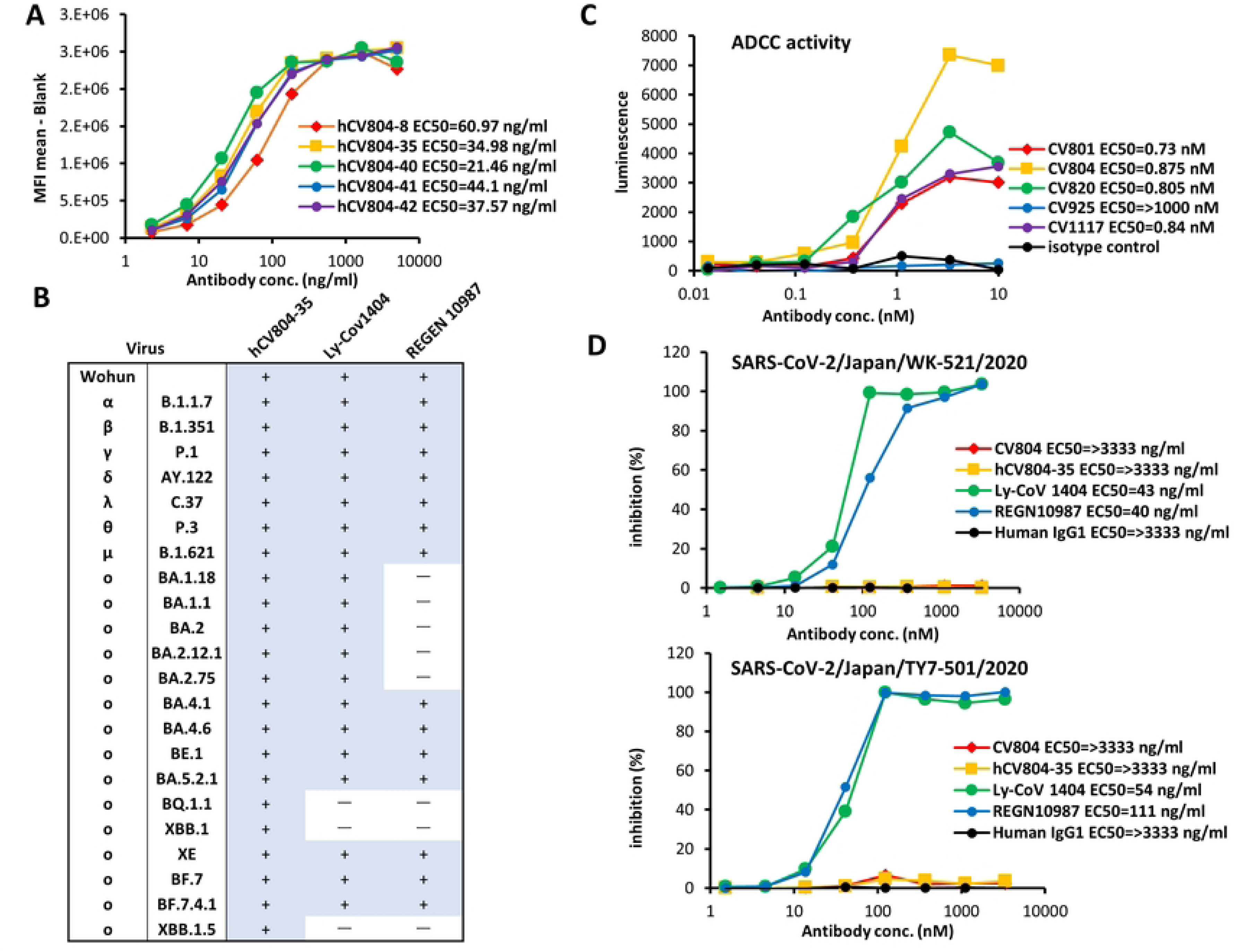
CV804 binds to SARS-CoV-2 and its variants, but does not neutralize them. (A) Five humanized CV804 antibodies were analyzed by flow cytometric assay of titrated candidates against SARS-CoV-2 spike protein expressing cells. Four parameter logistic curve fitted with EC_50_ in parentheses. Normalized by secondary alone and saturation signal. Representative data of two independent experiments are shown. (B) Evaluation of hCV804-35 antibody, REGN10987, and Ly-CoV1404 antibody binding to cells infected with SARS-CoV-2 variants. “+” indicates positive binding, “-” indicates negative binding. (C) Evaluation of antibody-dependent cell-mediated cytotoxicity ADCC activity against spike-expressing cells for mouse derived antibodies (CV801, CV804, CV820, CV925, CV1117). Four out of the five antibodies (not CV925) were found to have ADC activity. The EC_50_ values for each antibody indicate that the ADC activity was below the detectable limit and therefore the value could not be determined. (D) In vitro cell infection inhibition experiments were conducted. The viral strains used were hCoV-19/Japan/TY/WK-521/2020 (top) and SARS-CoV-2/Japan/TY-501/2020 (bottom), with VeroE6/TMPRSS2 cells being used as the target cells. The hCV804-35 antibody, REGN10987, and Ly-CoV1404 were measured in this experiment. The antibody concentration that inhibited cell death by 50% was defined as the IC_50_. Both REGN10987 and Ly-CoV1404 exhibited strong in vitro antiviral activity, whereas the CV804 antibody from the present study did not show any in vitro antiviral activity even at high concentrations.

To gain deeper insight into the characteristics of CV804, we assessed its binding to spike protein-expressing cells after infection using various mutant strains, including the Omicron variant. Unlike S1 antibodies such as REGN10987 (imdevimab)^21^ and LY-CoV1404 (bebtelovimab)^22^, CV804 demonstrated binding activity against multiple strains, including B.1.1.7 (alpha), B.1.351 (beta), P.1 (gamma), B.1.617.1 (kappa), B.1.617.2 (delta), and B.1.1.529 (omicron, BA.1), as well as the Omicron subvariants (BA.2.75, BA.4.1, BA.4.6, BE.1, BA.5.2.1, BQ.1.1, XBB.1, XE, BF.7, BF.7.4.1) (Fig 2B). It displayed remarkable resilience against viral mutations. We also evaluated the ADCC activity of CV804 and found that it had robust activity (Fig 2C). However, CV804 did not exhibit inhibitory activity against SARS-CoV-2 in a virus neutralization test using VeroE6/TMPRSS2 cells (Fig 2D). Furthermore, our investigation did not detect any membrane fusion-inhibiting activity of CV804 in a cell fusion induction assay system utilizing spike-expressing cells (Fig 2E).

### Antiviral activity of non-neutralizing CV804 against SARS-CoV-2

To evaluate the therapeutic effects of CV804 in vivo, female BALB/c mice were intranasally infected with 1.0×10^5^ TCID_50_/mouse of SARS-CoV-2 hCoV-19/Japan/TY7-501/2021 (gamma strain). Immediately after infection, the mice were intravenously administered 40 mg/kg of CV804 or isotype control. All mice were observed daily for survival and body weight changes (Figs 3B and 3C). In the group treated with 40 mg/kg of CV804, the body weight of the mice decreased by approximately 15% until day 4 and recovered to the level comparable to uninfected mice by day 6 (Fig 3B). In the isotype-treated group, all mice reached the humane endpoint or died by day 5, whereas 80% of the mice in the CV804-treated group survived, indicating an improvement in survival rate due to CV804 administration (Fig 3C). This effect was diminished by using the LALA mutant, which eliminates the Fc effector function of IgG, suggesting the possible contribution of ADCC activity to the therapeutic efficacy. However, the improvement in disease pathology observed with CV804 administration in this mouse model was limited. Although some degree of improvement was observed, the therapeutic effect was inferior to that of the neutralizing antibody (REGN10987) used as a control drug.

**Fig 3.**
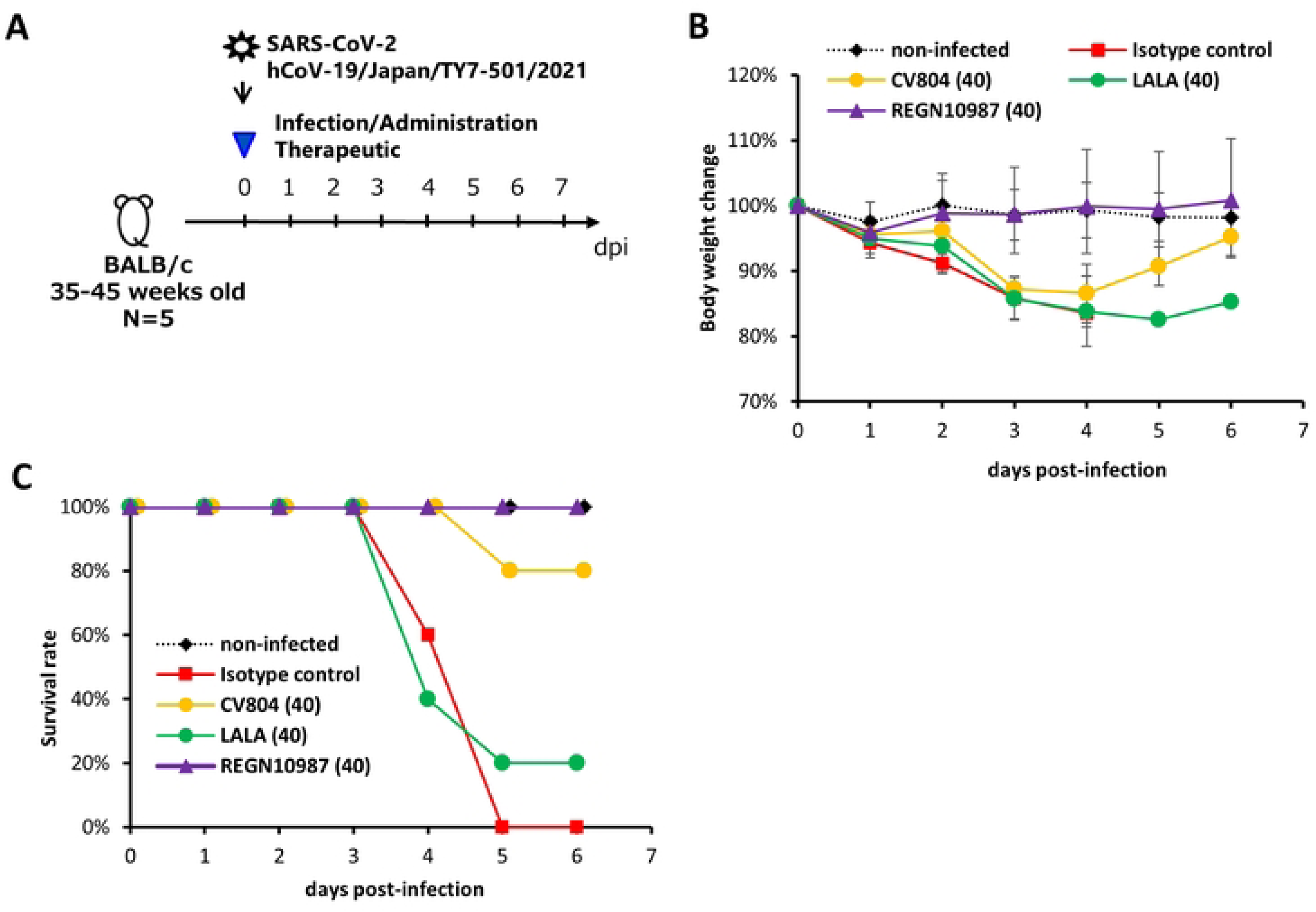
Antiviral activity of non-neutralizing CV804 against SARS-CoV-2. The therapeutic effects in aged mice were evaluated. (A) Outline for treatment protocol. Mice were intranasally inoculated with 50 μL of hCoV-19/Japan/TY7-501/2021 (1.00x10^5^ TCID50) under anesthesia. Non-infected control mice were intranasally inoculated with 50 μL of vehicle under anesthesia. Starting from immediately after virus infection, mice (n = 5/group) were intravenously administered a single dose of 40 mg/kg of CV804, CV804-LALA, and REGN10987, or a single dose of 40 mg/kg of C1.18.4 (isotype control). The body weight (B) and survival rate (C) of the mice were assessed once daily until day 6 post-virus infection.

### The epitope of CV804 is buried within the prefusion spike trimer

To further validate the conservation of the epitope recognized by CV804 and investigate the molecular basis of its broad specificity, antigen binding, and epitope interaction, we generated a spike protein ectodomain trimer and performed HDX-MS analysis to identify the CV804 epitope. We detected a peptide region within the CV804 binding trimer that showed more than 5% decrease in deuterium exchange efficiency compared to the isolated trimer, indicating the core of the epitope. We confirmed the binding of CV804 and humanized CV804 to the spike protein at positions 960-1001 (SARS-CoV-2 numbering) (Fig 4A). The data obtained as described above revealed the epitope region of CV804 and the specificity of the cross-reactivity of the antibody. Based on the reported higher order structure of the spike protein, we considered the rational relationship between the epitope and cross-reactivity.

**Fig 4.**
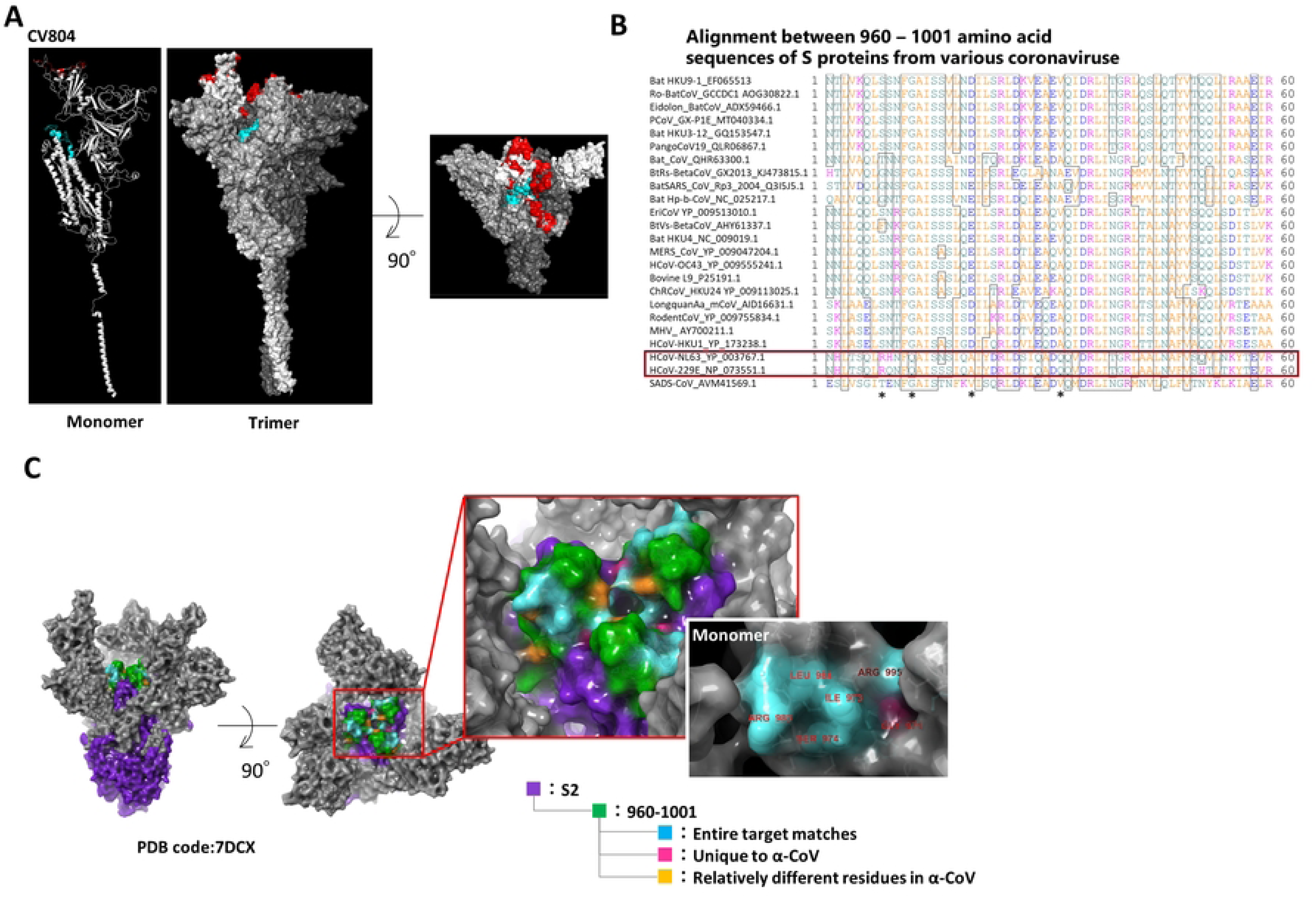
The epitope of CV804 is buried within the prefusion spike trimer. We performed epitope mapping of the CV804 antibody binding to the spike protein of SARS-CoV-2 using HDX-MS. (A) The regions detected by HDX-MS were plotted onto the three-dimensional structure (PDB ID: 6vsb_1_1_1) shown in red for the ACE2 binding recognition domain and in cyan for the regions detected by HDX-MS. The figures from left to right depict the monomer and trimer, with the figure on the right showing a top view of the spike trimer. It is speculated that the epitope region of CV804 is exposed due to structural changes when it adopts the UP form, similar to the class4 antibody epitope. (B) An amino acid sequence alignment of the spike protein epitope candidate region (960-1001) was performed. The red box represents alpha coronaviruses, and the asterisks indicate amino acids that are relatively different in alpha coronaviruses. (C) Highly conserved regions in alpha and beta coronaviruses were extracted and mapped onto the 2-up structure of the Wuhan strain crystal structure (PDB code: 7DCX). Purple represents the spike 2 protein, green represents the detected epitopes, cyan represents the regions that are conserved among all target coronaviruses within the detected epitopes, pink represents residues that are unique to alpha coronaviruses, and yellow represents amino acid residues that are relatively different in alpha coronaviruses. The red box enlarges the central part of the trimer. The white box illustrates the monomeric state of the region indicated by the red box, with the amino acid residue numbers indicating the universally conserved regions among coronaviruses and exposed residues on the surface of S2 protein.

For the epitope identified from the HDX-MS analysis, we performed primary sequence alignment of the spike proteins from 27 related coronaviruses (Fig 4B). We identified amino acid residues that are common to all virus species as well as those that are characteristic to beta coronaviruses but differ from alpha coronaviruses, and mapped them onto the reported higher order structure of CoV2 Spike 2-up form (PDB Code: 6BSV). As a result, many of the conserved amino acids in beta coronaviruses, which are included in the epitope region of CV804, were found to be located in exposed positions in the up form structure of the spike, strongly suggesting that the higher order structure composed of these residues is important for the binding of CV804 antibody (Fig 4C). In the alanine scan analysis using these mutants, the binding of CV804 antibody was abolished by mutations to alanine at residues L962, E988, Q992, L996, R1000, L1001, L1004, Y1007, and I1018, suggesting that these conserved residues are particularly important for the binding of CV804 to related coronaviruses (Table 4). In contrast, there were residues characteristic of alpha coronaviruses in the vicinity of these conserved residues, indicating a possible reason for the lack of cross-reactivity of CV804 with alpha coronaviruses. It should be noted that alanine mutations at these residues did not affect the reactivity of CV804, suggesting that the amino acid residues do not directly participate in the antigen-antibody interaction and are predicted to tolerate point mutations.

**Table 4.** Epitope mapping of the CV804 antibody against the spike protein of SARS-CoV-2 using alanine scanning.

| Ref<br>residue | Position | Variant<br>residue | CV804 |
| --- | --- | --- | --- |
| N | 960 | A | + |
| T | 961 | A | + |
| L | 962 | A | — |
| V | 963 | A | + |
| K | 964 | A | + |
| Q | 965 | A | + |
| L | 966 | A | + |
| S | 967 | A | + |
| S | 968 | A | + |
| F | 970 | A | + |
| I | 973 | A | + |
| S | 974 | A | + |
| N | 978 | A | + |
| D | 979 | A | + |
| I | 980 | A | + |
| L | 981 | A | + |
| R | 983 | A | + |
| L | 984 | A | + |
| D | 985 | A | + |
| K | 986 | A | + |
| V | 987 | A | + |
| E | 988 | A | — |
| E | 990 | A | + |
| V | 991 | A | + |
| Q | 992 | A | — |
| I | 993 | A | + |
| D | 994 | A | + |
| R | 995 | A | + |
| L | 996 | A | — |
| I | 997 | A | + |
| T | 998 | A | + |
| G | 999 | A | + |
| R | 1000 | A | — |
| L | 1001 | A | — |
| Q | 1002 | A | + |
| S | 1003 | A | + |
| L | 1004 | A | — |
| Q | 1005 | A | + |
| T | 1006 | A | + |
| Y | 1007 | A | — |
| V | 1008 | A | + |
| T | 1009 | A | + |
| Q | 1010 | A | + |
| Q | 1011 | A | + |
| L | 1012 | A | + |
| I | 1013 | A | + |
| R | 1014 | A | + |
| E | 1017 | A | + |
| I | 1018 | A | — |
| R | 1019 | A | + |

## Discussion

Antibody therapeutics against novel coronavirus infection should possess excellent specificity and efficacy. However, problems arise due to the emergence of virus variants resistant to nAbs, resulting in loss of efficacy, as well as the difficulty in generating broadly reactive antibodies that can target diverse viruses. In this study, we successfully developed an anti-SARS-CoV-2 spike 2 antibody CV804 that exhibits therapeutic effects by utilizing antibody effector functions in the host, which is distinct from traditional virus-nAbs. Despite lacking virus-neutralizing activity, CV804 demonstrated a certain degree of improvement in disease pathology in a lethal infection mouse model. Furthermore, CV804 recognizes a unique epitope that is different from other antibody therapeutics targeting the critical site RBD in virus-infected individuals, thereby specializing in supporting the host immune response and inducing therapeutic effects.

The location of the epitope core of the CV804 antibody, identified by HDX-MS, encompasses approximately 40 amino acids, including a portion of the HR1 region in the S2 domain and a portion of the central helical domain in the primary structure of the spike. In the quaternary structure, this region is located at the S1 end of the trimeric S2 domain of the viral spike, surrounded by the S1 receptor-binding domain. The spike protein of SARS-CoV-2 adopts an equilibrium between the “down” and “up” conformations, with the receptor-binding domain in the S1 domain binding to the ACE2 receptor on the surface of human cells, stabilizing the “up” conformation. In this “up” conformation, the S1 domain is distanced from the S2 domain, exposing the region of the S2 domain responsible for membrane fusion required for viral entry. The epitope structure of CV804 is located within the S2 region but is covered by the S1 domain and positioned inside the trimeric spike structure surrounded by the S1 receptor-binding domain in the “down” conformation. However, it becomes exposed when the spike protein adopts the “up” conformation. Due to the strong reactivity of CV804 towards cells expressing the spike protein, it is speculated that the exposed higher-order structure in the “up” conformation of the spike is present on the spike protein of infected cells. On the other hand, since CV804 does not exhibit in vitro infection-blocking activity (neutralizing activity), it is considered that the binding of the antibody to this epitope region does not possess inhibitory activity against the membrane fusion activity required for virus infection, which involves the S2 domain.

The unique epitope structure of the CV804 antibody is a three-dimensional epitope on the spike protein that is shared among many related coronaviruses. CV804 exhibits a high degree of cross-reactivity with various mutant strains and related coronaviruses. Structural data indicate that several key residues involved in the antibody-antigen interaction are either completely or partially hidden within the pre-fusion S trimer. The loss of neutralizing capability of CV804 may be due to limited epitope exposure. This suggests that structural changes occur during the transition from the pre-fusion state to the post-fusion state, leading to increased epitope exposure and enabling binding of CV804.

The S2 region of coronaviruses is highly conserved and represents a promising target for a wide range of coronavirus antibodies and vaccines, compared to the S1 region^17,23–25^. Several nAbs such as S2P6, CC40.8, IgG22, B6, 28D9, and CV3-25 have been isolated from convalescent patients, vaccinated individuals, and mice^13,14,26–28^. Consistent with the highly conserved S2 stem helix epitope among beta-coronaviruses, these nAbs typically demonstrate a broad neutralizing breadth against beta-coronaviruses, including SARS-CoV-1 and SARS-CoV-2^13,14^. In our study as well, CV804 exhibited extensive reactivity against beta-coronaviruses, similar to or surpassing the aforementioned antibodies. The highly conserved epitope recognized by CV804 leads to its extensive reactivity. Mutations within the epitope residues of CV804 in SARS-CoV-2 sequences were found to be less than 0.032%, and all relevant SARS-CoV-2 variants do not have residue mutations within this epitope. Notably, major natural variations within the CV804 epitope did not affect its binding activity. Some mutations can significantly reduce viral infectivity, thus potentially limiting the spread of the virus. We concluded that CV804, with its broad spectrum, represents an attractive means to address viral escape mutations and should be a valuable tool in tackling future outbreaks of novel coronaviruses.

We discovered that the binding of ACE2 promotes the exposure of the S2’ site and fusion peptide, possibly removing the obstacle posed by this epitope through structural reorganization of the S protein and providing access to CV804. Additionally, CV804 antibody targets the spike protein expressed on infected cells and is specialized in antibody effector functions in the body, and thus does not require virus neutralizing activity. It binds to the less mutable S2 region to impair infected cells, demonstrating efficacy in animal experiments. CV804 synergizes with human ACE2 to prevent SARS-CoV-2 infection, and we speculate that the binding of CV804 to the S structure, which includes the S2’ site and fusion peptide, may have synergistic effects with receptor priming or receptor blocking processes and ADCC activity. Our data also suggest that some RBD antibodies may provide possible combinations with CV804 and human ACE2 or RBD antibodies for the prevention or treatment of SARS-CoV-2 infection. However, the mechanism of action of CV804 supports the immune response mediated by the host and therefore, monitoring the immune function of model mice is necessary, but the details are still unclear. The efficacy potential of CV804 as a monotherapy is considered to be insufficient compared to previous nAbs. It is important to elucidate the epitope details of the CV804 antibody and understand the antibody characteristics, such as cross-reactivity, for future development of broad-spectrum anti-coronavirus antibody therapeutics and further research on vaccine development to efficiently induce useful antibody responses.

We conducted binding evaluation on the expression cells of spike proteins belonging to the genus of coronaviruses. The five obtained antibodies, CV801, CV804, CV820, CV925, and CV1117, reacted with more than 52% of the different spike proteins among the 17 different viral spikes examined. CV804 reacted with more than 70% of the different spike proteins among the 17 different viral spikes. “+”, “-”, and “N.T.” represent positive binding, negative binding, and experiment not performed, respectively.

To confirm the reactivity of the antibodies of the present study to various SARS-CoV-2 spike protein variants, including SARS-CoV-2 wild-type (Genbank ACC No. QHD43416.1), alpha variant (GISAID ID: EPI_ISL_768526), gamma variant (GISAID ID: EPI_ISL_833366), and omicron variant (GISAID ID: EPI_ISL_6640917), the experiments were conducted using flow cytometry. The cells expressing each full-length spike protein were incubated with the mouse antibodies. The amino acid variants of the SARS-CoV-2 spike protein as shown in Table 4 represent the mutation of the amino acid residue at position 452 from L to R in the wild-type SARS-CoV-2 spike protein (Genbank ACC No. QHD43416.1), with other variants having similar mutations. Amino acid positions 686 and above represent mutations found in the extracellular domain of the S2 subunit, which are present in any of the naturally occurring SARS-CoV-2 spike protein variants. In the table, “+”, “-”, and “N.T.” represent positive binding, negative binding, and experiment not performed, respectively. It was found that the antibodies of the present study bind to all tested variants.

To determine the binding sites (epitopes) of the CV804 antibody from the amino acid sequence of the SARS-CoV-2 spike protein (Genbank ACC No. QHD43416.1), mass spectrometry was performed using hydrogen/deuterium exchange epitope mapping. The hydrogen-deuterium exchange rates of each peptide were automatically calculated from the mass spectra obtained in the full-scan mode in the deuterium exchange experiment. Peptides with calculated hydrogen-deuterium exchange rates were filtered according to the criteria of a difference (ΔD%) in hydrogen-deuterium exchange rates between samples with and without the presence of hCV804-40 antibody in two or more adjacent peptides being 10% or more. The amino acid residues of the SARS-CoV-2 spike protein corresponding to the region containing peptides that met the criteria were identified as the epitope of the hCV804-40 antibody. The sequence coverage of the target protein was 98.5%, and a total of 737 peptides were identified in the samples with and without the presence of the hCV804-40 antibody. Among the identified 737 peptides, 36 peptides met the criteria of a difference (ΔD%) in hydrogen-deuterium exchange rates between samples with and without the presence of the hCV804-40 antibody being 10% or more. These 36 peptides were found to contain 42 amino acid residues (sequence number 49) corresponding to amino acid residues 960-1001 (sequence number 48) of the SARS-CoV-2 spike protein (Genbank ACC No. QHD43416.1). Therefore, amino acid residues of the SARS-CoV-2 spike protein to which the hCV804-40 antibody binds are located at positions 960-1001. The table data show the sequence numbers, amino acid sequences, positions of amino acid residues in the SARS-CoV-2 spike protein, and ΔD% for the above-mentioned 36 peptides. Similar experiments were conducted for the CV804 antibody, and similar results were obtained, indicating that amino acid residues 960-1001 of the SARS-CoV-2 spike protein are also included in the epitope.

50 site-specific mutants were created by replacing amino acids from position 960 to 1018 of the SARS-CoV-2 spike protein with alanine, and the reactivity of the CV804 antibody was evaluated. The results are represented as “+” for positive binding and “-” for negative binding. The overall structure of the protein remains intact by substituting each amino acid with alanine, where each amino acid residue is replaced with a methyl group that forms alanine, allowing detection of the function of the original amino acid residue before substitution, which is necessary for the antigen-antibody interaction. Alanine is used because it has a small size and a chemically inactive methyl group, which can mimic the secondary structure shown by many other amino acids. The results revealed that mutants L962A, E988A, Q992A, L996A, R1000A, L1001A, L1004A, Y1007A, and I1018A showed negative binding reactivity with the CV804 antibody. The amino acids replaced by alanine in these mutants are important amino acids for the binding of the CV804 antibody and constitute the epitope of the CV804 antibody. Therefore, the loss of reactivity of the CV804 antibody indicates that the region from position 962 to 1018 of the spike protein of SARS-CoV-2 is included in the epitope sequence of the CV804 antibody.

## Acknowledgements

We are grateful to all our colleagues who participated in the COVID-19 antiviral program at Shionogi and thank Shionogi TechnoAdvance Research Co., Ltd. for technical support in the pharmacological studies. The authors thank Sayuri Okamoto, Mayumi Niiyama, Kayoko Kato, Akiko Okabe, Madoka Takeshima, Eiko Moriishi, and Futaba Makimura who participated in the COVID-19 antiviral program at Laboratory of Antibody Design, Center for Drug Design Research, National Institutes of Biomedical Innovation, Health and Nutrition for technical support in the pharmacological studies.

## Supporting information

**S1 Table. The numbers of euthanized, died or survived mice in the study for examining the lethality in SARS-CoV-2 infected mice.**

**S1 Fig. Antibody Isolation and binding activity to S protein.** (A) Reactivity of CV804 antibody to SARS-CoV-2 spike protein variants expressed in cells. The following experiments were conducted to assess the reactivity of the present invention’s antibodies to amino acid variants of each SARS-CoV-2 spike protein. The table shows that the antibodies of the present study could bind to all tested variants. (B) Binding evaluation of CV804 antibody to cells expressing spike protein of related coronaviruses. Expression plasmid vectors for spike protein were generated, and the binding of CV804 antibody to transfected cell populations was assessed using the same method as in Table 1. CV804 antibody exhibited reactivity towards over 73% of the distinct spike proteins among the 15 different viral strains.“+” indicates positive binding, “-” indicates negative binding.

**S2 Fig. Evaluation of hCV804-35 antibody, REGN10987, and Ly-CoV1404 antibody binding to cells infected with SARS-CoV-2 variants.** To further understand the characteristics of CV804, we conducted a study on its binding to cells expressing spike protein post-infection using various mutant strains, including the Omicron variant. Unlike S1 antibodies such as REGN10987 and LY-CoV1404, CV804 exhibited binding activity against multiple strains, including B.1.1.7, B.1.351, P.1, B.1.617.1, B.1.617.2, and B.1.1.529 (BA.1), as well as the Omicron subvariants (BA.2.75, BA.4.1, BA.4.6, BE.1, BA.5.2.1, BQ.1.1, XBB.1, XE, BF.7, BF.7.4.1). The hCV804-40 antibody demonstrated spot confirmation for all strains. However, Ly-Cov1404 and REGEN 10987 did not show spot confirmation for three to four types of οmicron strains.

## References

1. Onyeaka, H., Anumudu, C. K., Al-Sharify, Z. T., Egele-Godswill, E. & Mbaegbu, P. COVID-19 pandemic: A review of the global lockdown and its far-reaching effects. Science Progress vol. 104 Preprint at 10.1177/00368504211019854 (2021).

2. Carstens, E. B. Ratification vote on taxonomic proposals to the International Committee on Taxonomy of Viruses (2009). Arch Virol 155, (2010).

3. 3. Chen, Y., Liu, Q. & Guo, D. Emerging coronaviruses: Genome structure, replication, and pathogenesis. Journal of Medical Virology vol. 92 Preprint at 10.1002/jmv.25681 (2020).

4. 4. Paules, C. I., Marston, H. D. & Fauci, A. S. Coronavirus Infections-More Than Just the Common Cold. JAMA - Journal of the American Medical Association vol. 323 Preprint at 10.1001/jama.2020.0757 (2020).

5. 5. Cui, J., Li, F. & Shi, Z. L. Origin and evolution of pathogenic coronaviruses. Nature Reviews Microbiology vol. 17 Preprint at 10.1038/s41579-018-0118-9 (2019).

6. 6. Telenti, A., et al. After the pandemic: perspectives on the future trajectory of COVID-19. Nature vol. 596 Preprint at 10.1038/s41586-021-03792-w (2021).

7. Weisblum, Y. et al. Escape from neutralizing antibodies 1 by SARS-CoV-2 spike protein variants. Elife 9, (2020).

8. Suryadevara, N. et al. Neutralizing and protective human monoclonal antibodies recognizing the N-terminal domain of the SARS-CoV-2 spike protein. Cell 184, (2021).

9. Yi, C. et al. Key residues of the receptor binding motif in the spike protein of SARS-CoV-2 that interact with ACE2 and neutralizing antibodies. Cell Mol Immunol 17, (2020).

10. Yu, F. et al. Receptor-binding domain-specific human neutralizing monoclonal antibodies against SARS-CoV and SARS-CoV-2. Signal Transduction and Targeted Therapy vol. 5 Preprint at 10.1038/s41392-020-00318-0 (2020).

11. Du, L., Yang, Y. & Zhang, X. Neutralizing antibodies for the prevention and treatment of COVID-19. Cellular and Molecular Immunology vol. 18 Preprint at 10.1038/s41423-021-00752-2 (2021).

12. Shi, R. et al. A human neutralizing antibody targets the receptor-binding site of SARS-CoV-2. Nature 584, (2020).

13. Zhou, P. et al. A human antibody reveals a conserved site on beta-coronavirus spike proteins and confers protection against SARS-CoV-2 infection. Sci Transl Med 14, (2022).

14. Pinto, D. et al. Broad betacoronavirus neutralization by a stem helix–specific human antibody. Science (1979) 373, (2021).

15. Wang, C. et al. A conserved immunogenic and vulnerable site on the coronavirus spike protein delineated by cross-reactive monoclonal antibodies. Nat Commun 12, (2021).

16. Shiakolas, A. R. et al. Cross-reactive coronavirus antibodies with diverse epitope specificities and Fc effector functions. Cell Rep Med 2, (2021).

17. Sun, X. et al. Neutralization mechanism of a human antibody with pan-coronavirus reactivity including SARS-CoV-2. Nat Microbiol 7, (2022).

18. 18. Vanderven, H. A., Jegaskanda, S., Wheatley, A. K. & Kent, S. J. Antibody-dependent cellular cytotoxicity and influenza virus. Current Opinion in Virology vol. 22 Preprint at 10.1016/j.coviro.2016.12.002 (2017).

19. Yu, Y. et al. Antibody-dependent cellular cytotoxicity response to SARS-CoV-2 in COVID-19 patients. Signal Transduct Target Ther 6, (2021).

20. 20. Harvey, W. T., et al. SARS-CoV-2 variants, spike mutations and immune escape. Nature Reviews Microbiology vol. 19 Preprint at 10.1038/s41579-021-00573-0 (2021).

21. Hansen, J. et al. Studies in humanized mice and convalescent humans yield a SARS-CoV-2 antibody cocktail. Science (1979) 369, (2020).

22. Westendorf, K. et al. LY-CoV1404 (bebtelovimab) potently neutralizes SARS-CoV-2 variants. Cell Rep 39, (2022).

23. Hu, B., Guo, H., Zhou, P. & Shi, Z. L. Characteristics of SARS-CoV-2 and COVID-19. Nature Reviews Microbiology vol. 19 Preprint at 10.1038/s41579-020-00459-7 (2021).

24. Guo, L. et al. Targetable elements in SARS-CoV-2 S2 subunit for the design of pan-coronavirus fusion inhibitors and vaccines. Signal Transduction and Targeted Therapy vol. 8 Preprint at 10.1038/s41392-023-01472-x (2023).

25. Silva, R. P. et al. Identification of a conserved S2 epitope present on spike proteins from all highly pathogenic coronaviruses. Elife 12, (2023).

26. Li, W. et al. Structural basis and mode of action for two broadly neutralizing antibodies against SARS-CoV-2 emerging variants of concern. Cell Rep 38, (2022).

27. Sauer, M. M. et al. Structural basis for broad coronavirus neutralization. Nat Struct Mol Biol 28, (2021).

28. Hsieh, C. L. et al. Stabilized coronavirus spike stem elicits a broadly protective antibody. Cell Rep 37, (2021).

29. Engen, J. R. & Smith, D. L. Investigating protein structure and dynamics by hydrogen exchange MS. Analytical Chemistry vol. 73 Preprint at 10.1021/ac012452f (2001).

30. Ehring, H. Hydrogen exchange/electrospray ionization mass spectrometry studies of structural features of proteins and protein/protein interactions. Anal Biochem 267, (1999).

